## Supplementary information for "Lifelong single-cell profiling of cranial neural crest diversification"

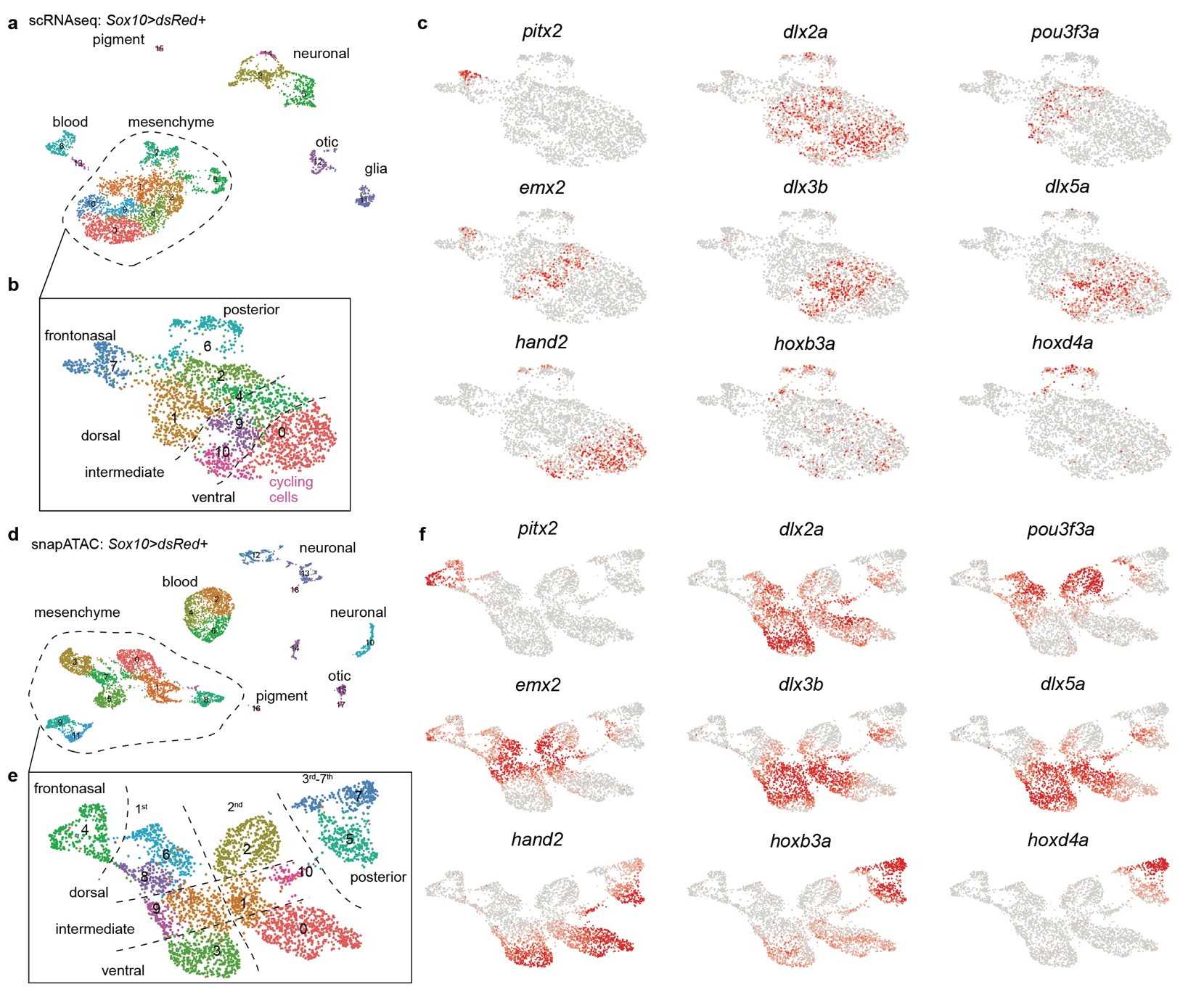
**Supplementary Figure 1. Single-cell datasets at 1.5 dpf. a,b**, scRNAseq UMAP for 1.5 dpf *Sox10>dsRed* dataset with box corresponding to ectomesenchyme subset (dashed outline). **c**, Feature plots of select genes for scRNAseq data. **d,e**, snATACseq UMAP for 1.5 dpf *Sox10>dsRed* dataset with box corresponding to ectomesenchyme subset (dashed outline). Arches are numbered at top in boxed region. **f**, Feature plots of select gene body activities for snATACseq data.

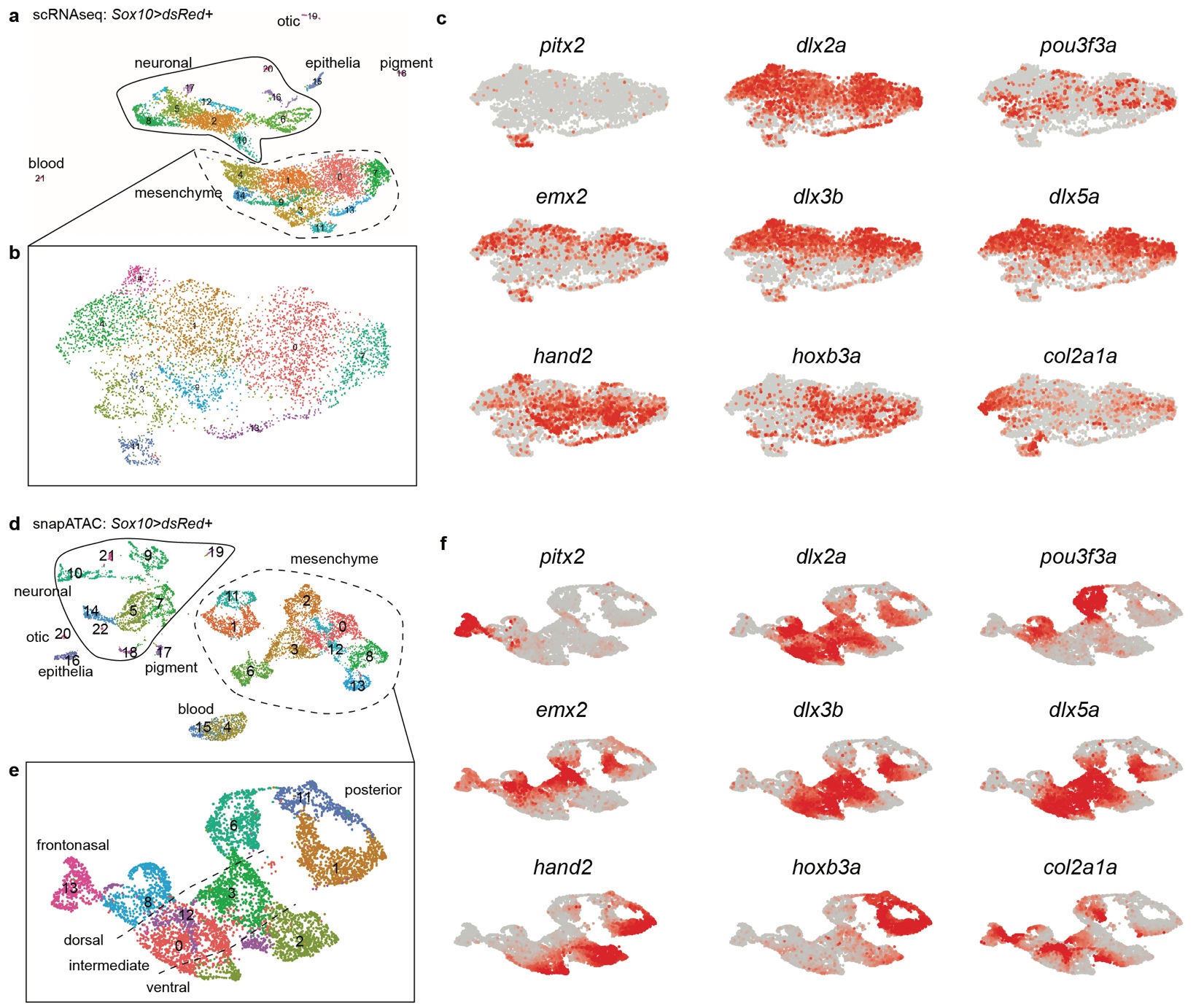

**Supplementary Figure 2. Single-cell datasets at 2 dpf.** **a,b**, scRNAseq UMAP for 2 dpf *Sox10>dsRed* dataset with box corresponding to ectomesenchyme subset (dashed outline). **c**, Feature plots of select genes for scRNAseq data. **d,e**, snATACseq UMAP for 2 dpf *Sox10>dsRed* dataset with box corresponding to ectomesenchyme subset (dashed outline). Arches are numbered at top in boxed region. **f**, Feature plots of select gene body activities for snATACseq data.

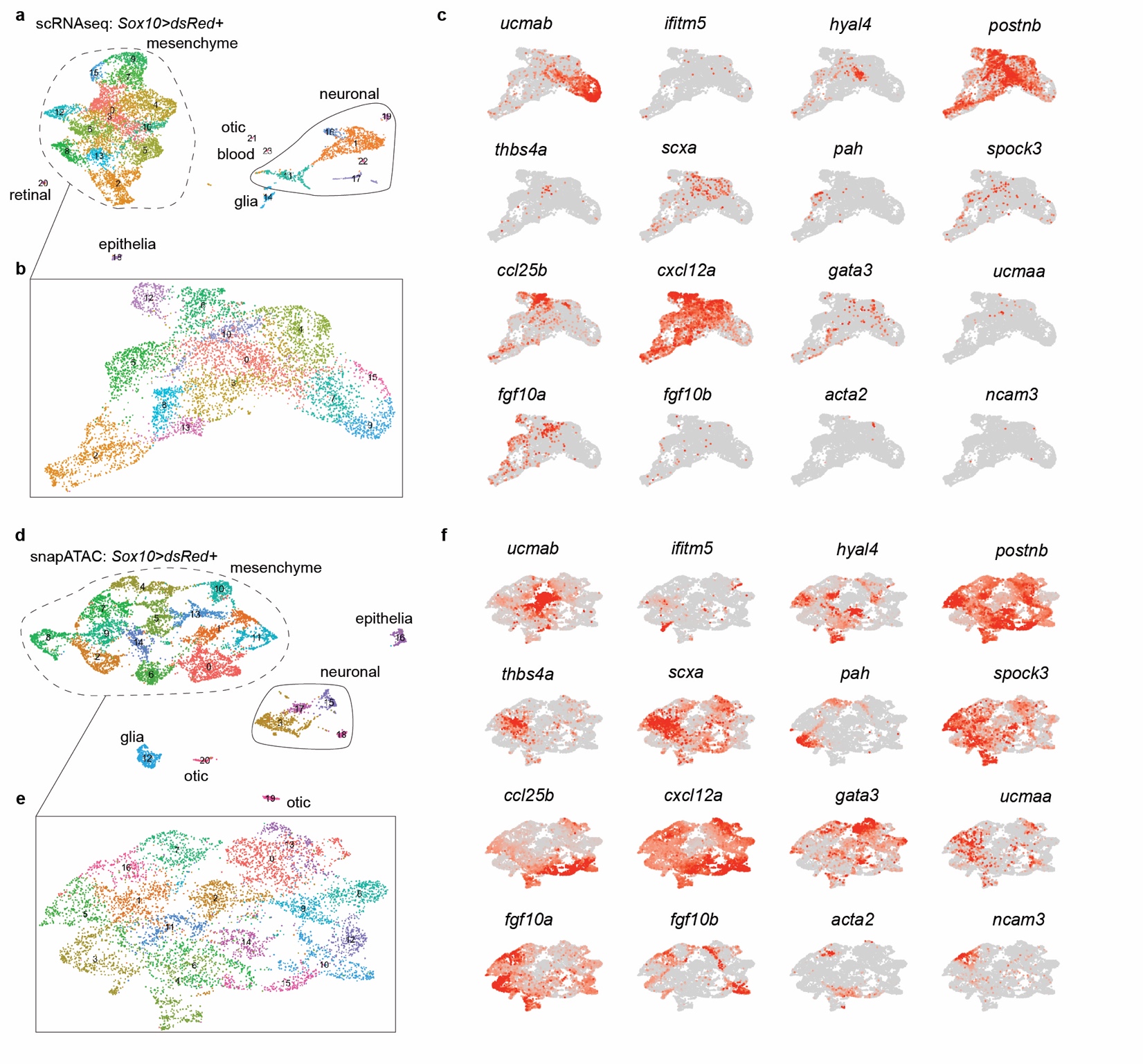
**Supplementary Figure 3. Single-cell datasets at 3 dpf.** **a,b**, scRNAseq UMAP for 3 dpf *Sox10>dsRed* dataset with box corresponding to ectomesenchyme subset (dashed outline). **c**, Feature plots of select genes for scRNAseq data. **d,e**, snATACseq UMAP for 3 dpf *Sox10>dsRed* dataset with box corresponding to ectomesenchyme subset (dashed outline). Arches are numbered at top in boxed region. **f**, Feature plots of select gene body activities for snATACseq data.

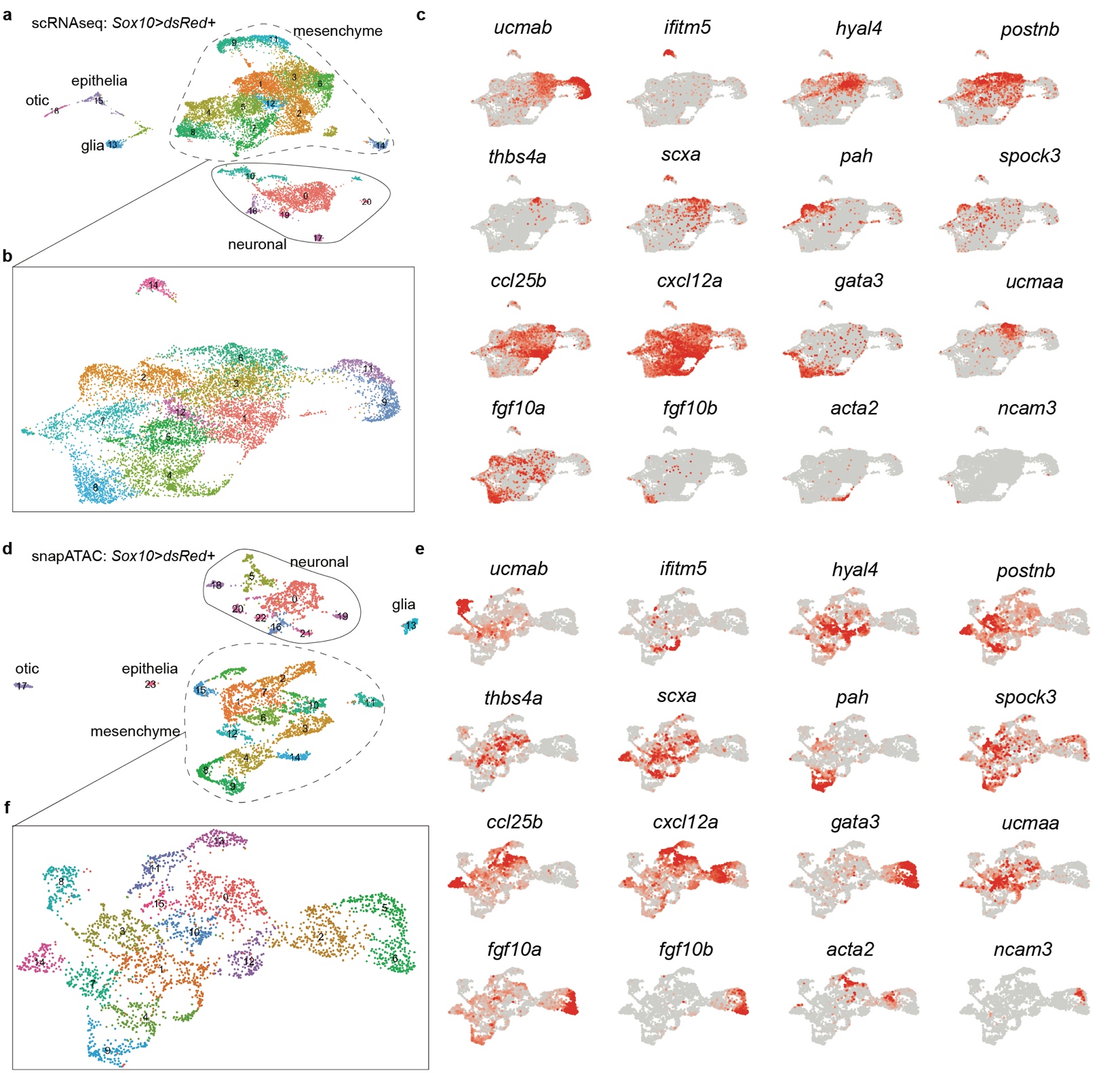

**Supplementary Figure 4. Single-cell datasets at 5 dpf. a,b**, scRNAseq UMAP for 5 dpf *Sox10>dsRed* dataset with box corresponding to ectomesenchyme subset (dashed outline). **c**, Feature plots of select genes for scRNAseq data. **d,e**, snATACseq UMAP for 5 dpf *Sox10>dsRed* dataset with box corresponding to ectomesenchyme subset (dashed outline). Arches are numbered at top in boxed region. **f**, Feature plots of select gene body activities for snATACseq data.

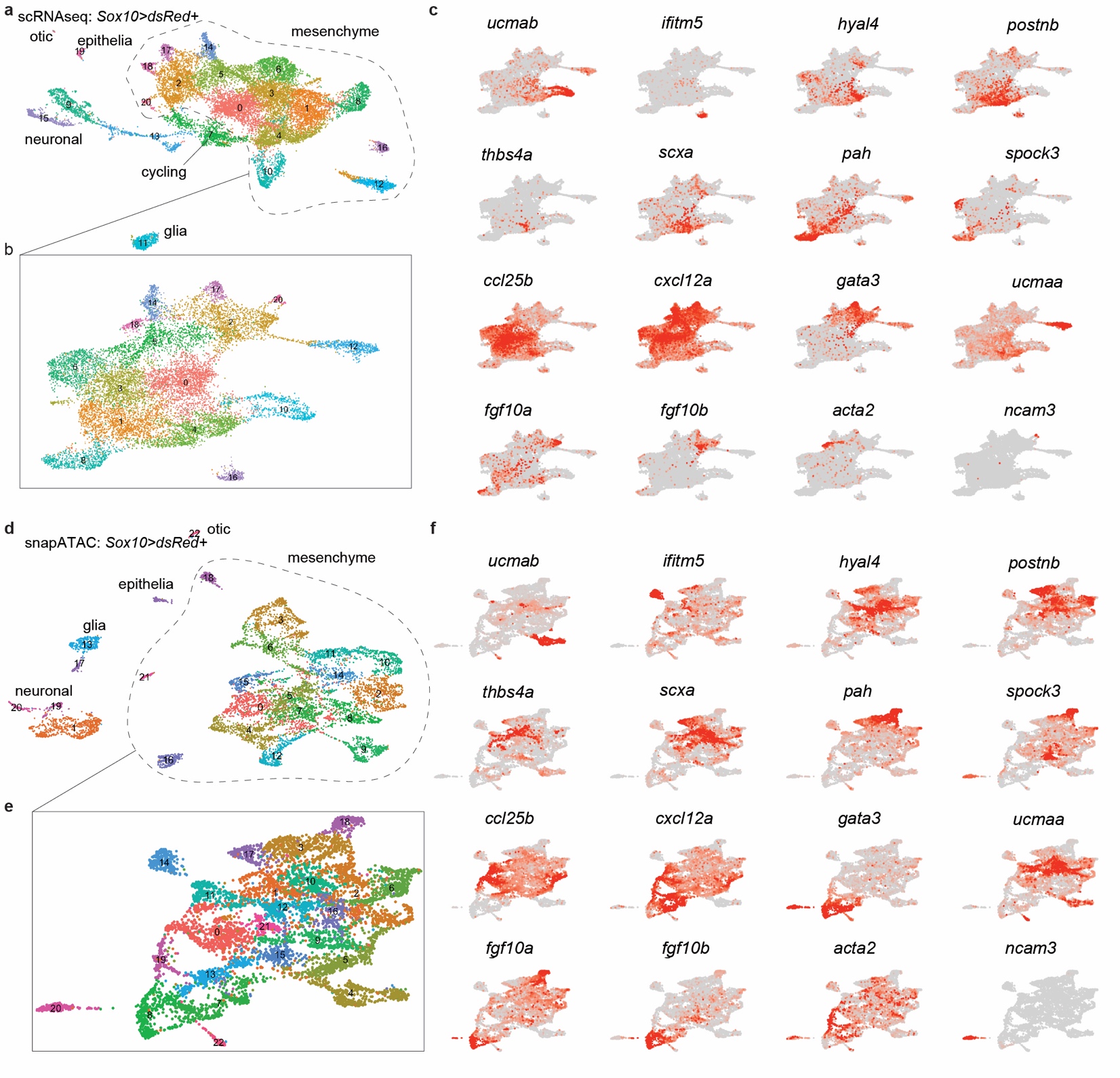
**Supplementary Figure 5. Single-cell datasets at 14 dpf. a,b**, scRNAseq UMAP for 14 dpf *Sox10>dsRed* dataset with box corresponding to ectomesenchyme subset (dashed outline). **c**, Feature plots of select genes for scRNAseq data. **d,e**, snATACseq UMAP for 14 dpf *Sox10>dsRed* dataset with box corresponding to ectomesenchyme subset (dashed outline). Arches are numbered at top in boxed region. **f**, Feature plots of select gene body activities for snATACseq data.

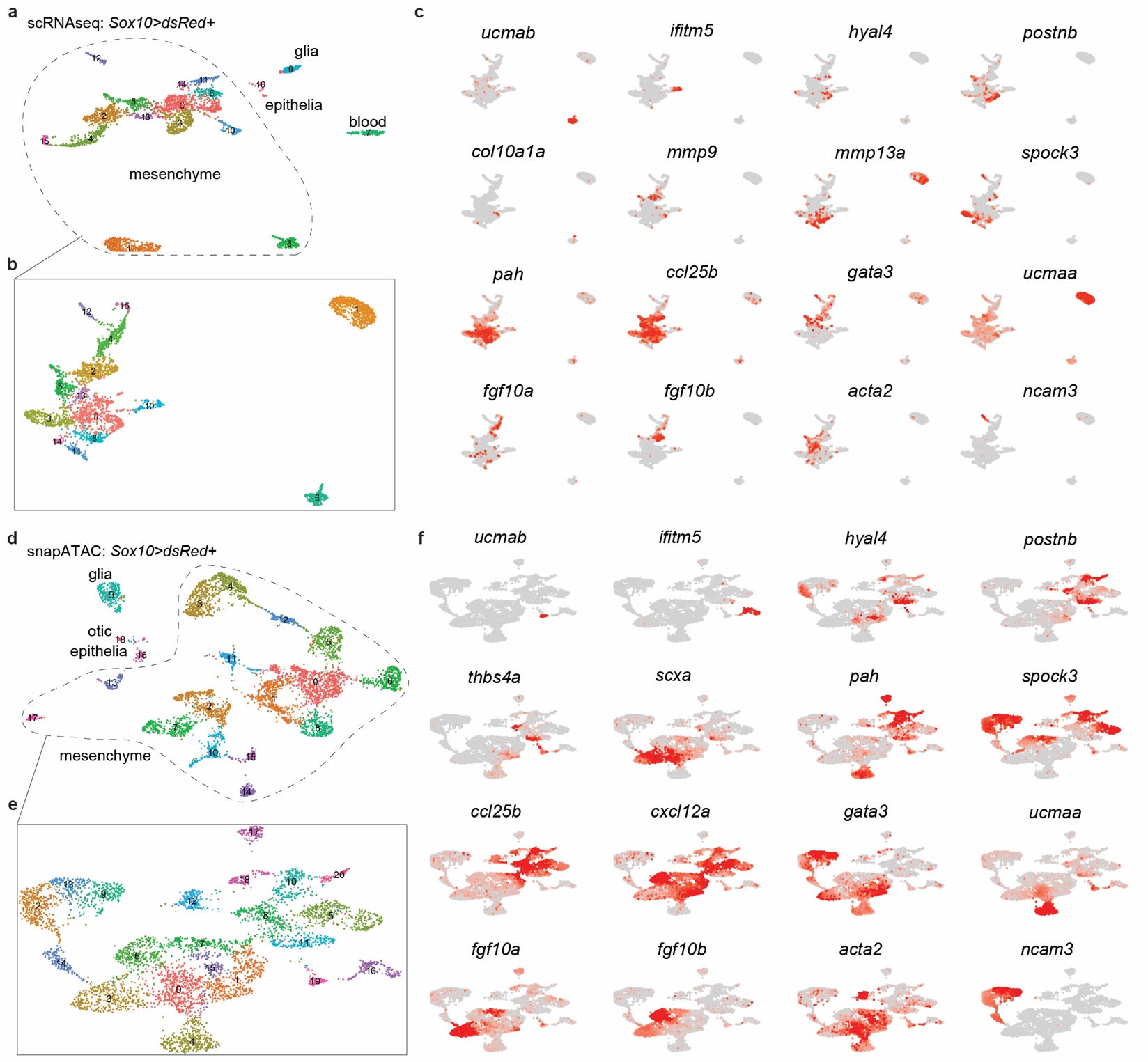

**Supplementary Figure 6. Single-cell datasets at 60 dpf. a,b**, scRNAseq UMAP for 60 dpf *Sox10>dsRed* dataset with box corresponding to ectomesenchyme subset (dashed outline). **c**, Feature plots of select genes for scRNAseq data. **d,e**, snATACseq UMAP for 60 dpf *Sox10>dsRed* dataset with box corresponding to ectomesenchyme subset (dashed outline). Arches are numbered at top in boxed region. **f**, Feature plots of select gene body activities for snATACseq data.

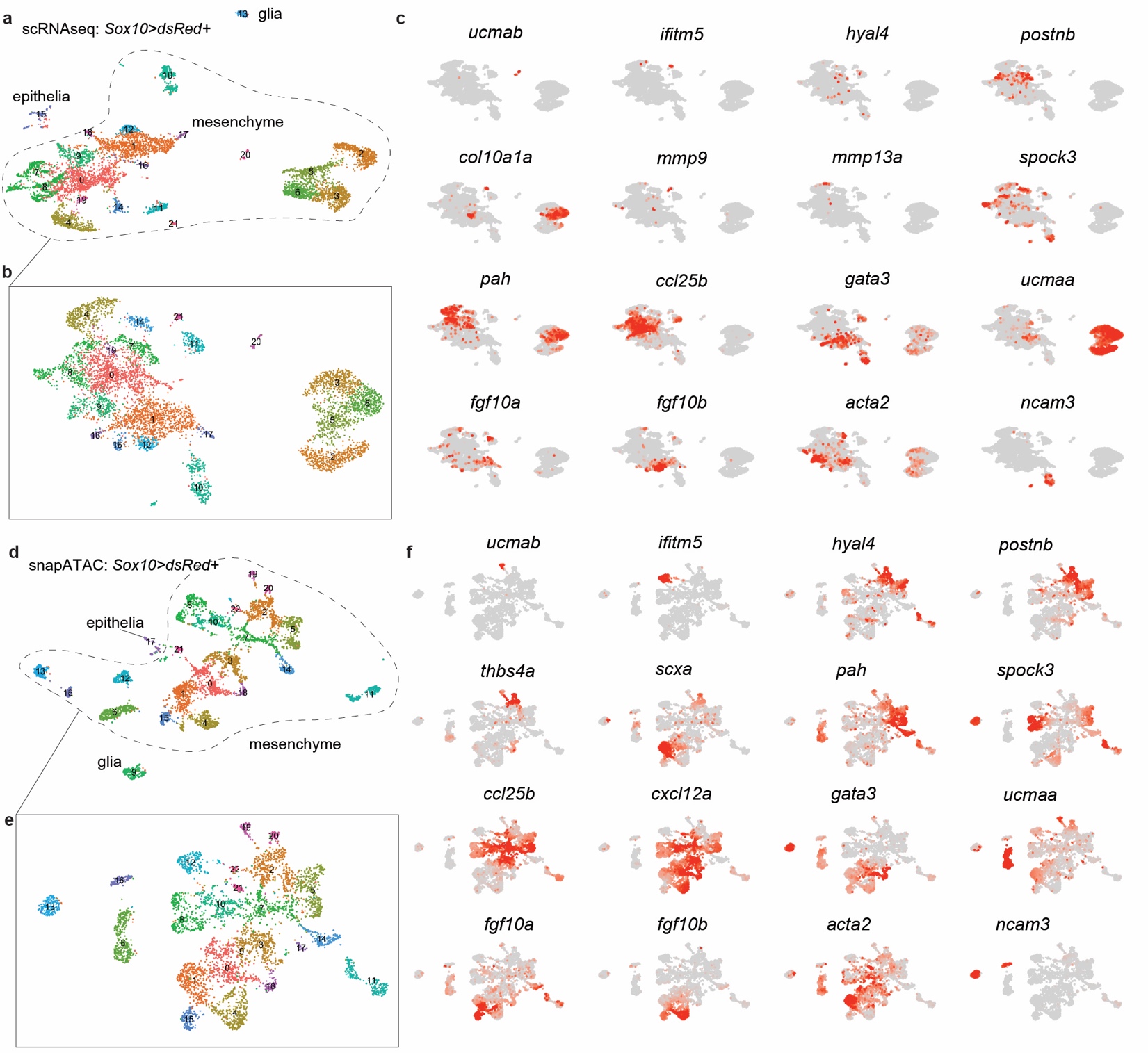

**Supplementary Figure 7. Single-cell datasets at adult stages.** **a,b**, scRNAseq UMAP for 150 dpf *Sox10>dsRed* dataset with box corresponding to ectomesenchyme subset (dashed outline). **c**, Feature plots of select genes for scRNAseq data. **d,e**, snATACseq UMAP for 210 dpf *Sox10>dsRed* dataset with box corresponding to ectomesenchyme subset (dashed outline). Arches are numbered at top in boxed region. **f**, Feature plots of select gene body activities for snATACseq data.

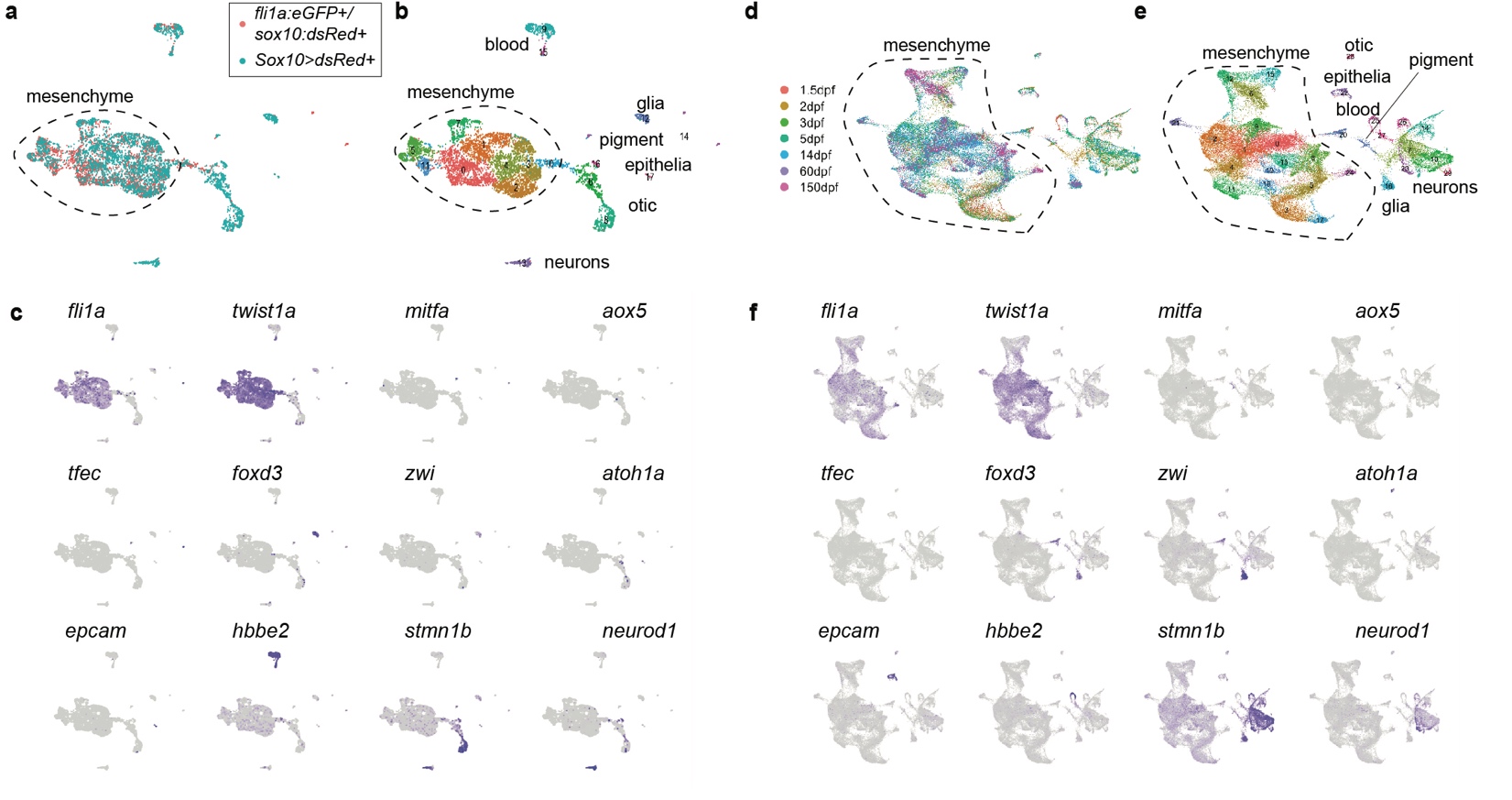

**Supplementary Figure 8. Single-cell analysis of ectomesenchyme and non-ectomesenchyme CNCC derivatives. a,** Co-clustering of *fli1a:eGFP+/sox10:dsRed+* and lineage-traced *Sox10>dsRed+* 1.5 dpf scRNAseq datasets using canonical correlation analysis (CCA) shows common contributions to ectomesenchyme (dashed outline) and blood (likely reflecting autofluorescence during sorting). As the *fli1a:eGFP* transgene is selective for ectomesenchyme CNCC derivatives, non-ectomesenchyme derivatives (glia, neurons, pigment cells) are only recovered in the *Sox10>dsRed+* dataset. Recovery of otic placode derivatives reflects *sox10* activity in this tissue, and a small number of epithelial cells are also selectively recovered in the *Sox10>dsRed+* dataset. **b**, Cell clusters in the combined datasets. **c**, Feature plots of select genes for ectomesenchyme (*fli1a*, *twist1a*), melanocytes (*mitfa*), xanthophores (*aox5*), iridiphores (*tfec*), Schwann cell glia (*foxd3*, *zwi*), otic cells (*atoh1a*), epithelia (*epcam*), blood (*hbbe2*), and neurons (*stmn1b*, *neurod1*). **d,e**, UMAP plot derived from CCA of all scRNAseq datasets (1.5, 2, 3, 5, 14, 60, 150 dpf) with colors showing stages (d) and clusters (e). **f**, Feature plots of select genes.

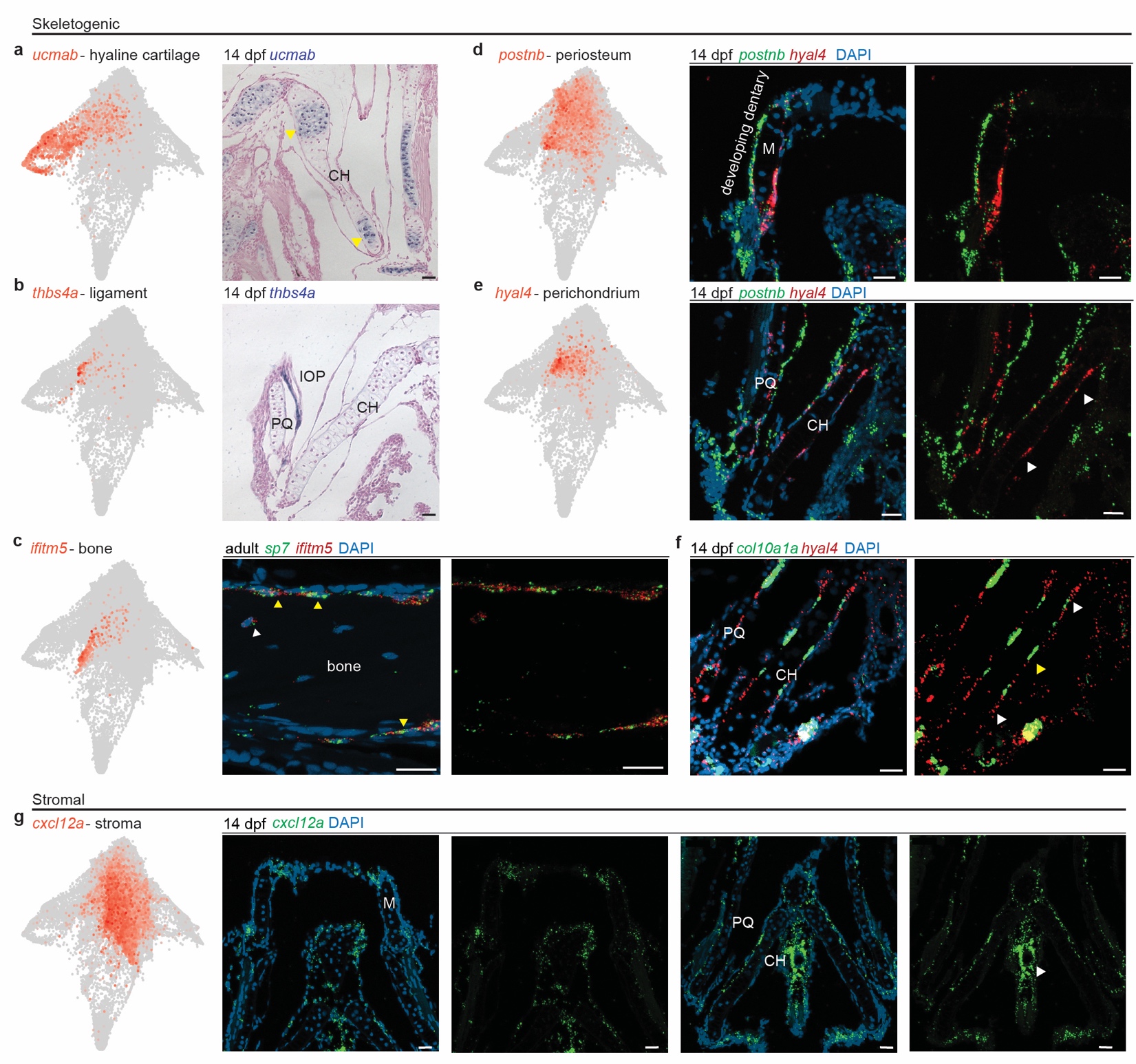

**Supplementary Figure 9. In vivo expression of markers for skeletogenic and stromal tissues. a,b**, STITCH plots show expression of *ucmab* and *thbs4a*. Colorimetric in situ hybridization on sections of the lower face/jaw region show expression of *ucmab* in cartilage, including growth plate chondrocytes (arrowheads) of the ceratohyal (CH), and expression of *thbs4a* in the interopercular-mandibular (IOP) ligament. **c-g**, STITCH plots and RNAscope in situ hybridization on sections of the lower face/jaw. In adult (49 dpf), *sp7* and *ifitm5* are co-localized in an embedded osteocyte (white arrowhead) and periosteal osteoblasts (yellow arrowheads) of adult bone. At 14 dpf, *hyal4* and *postnb* mark largely distinct populations surrounding cartilage, with *postnb* marking the developing dentary bone (consistent with periosteum identity). *hyal4* marks perichondrium surrounding ch growth plates (white arrowheads) in a largely complementary pattern to *col10a1a*+ osteoblasts in the periosteum (yellow arrowhead). At 14 dpf, *cxcl12a* is expressed broadly in mesenchyme, with particular enrichment around large diameter blood vessels (arrowhead) at the midline. DAPI labels nuclei in blue. M, Meckel’s cartilage; PQ, palatoquadrate. Scale bars = 20 um.

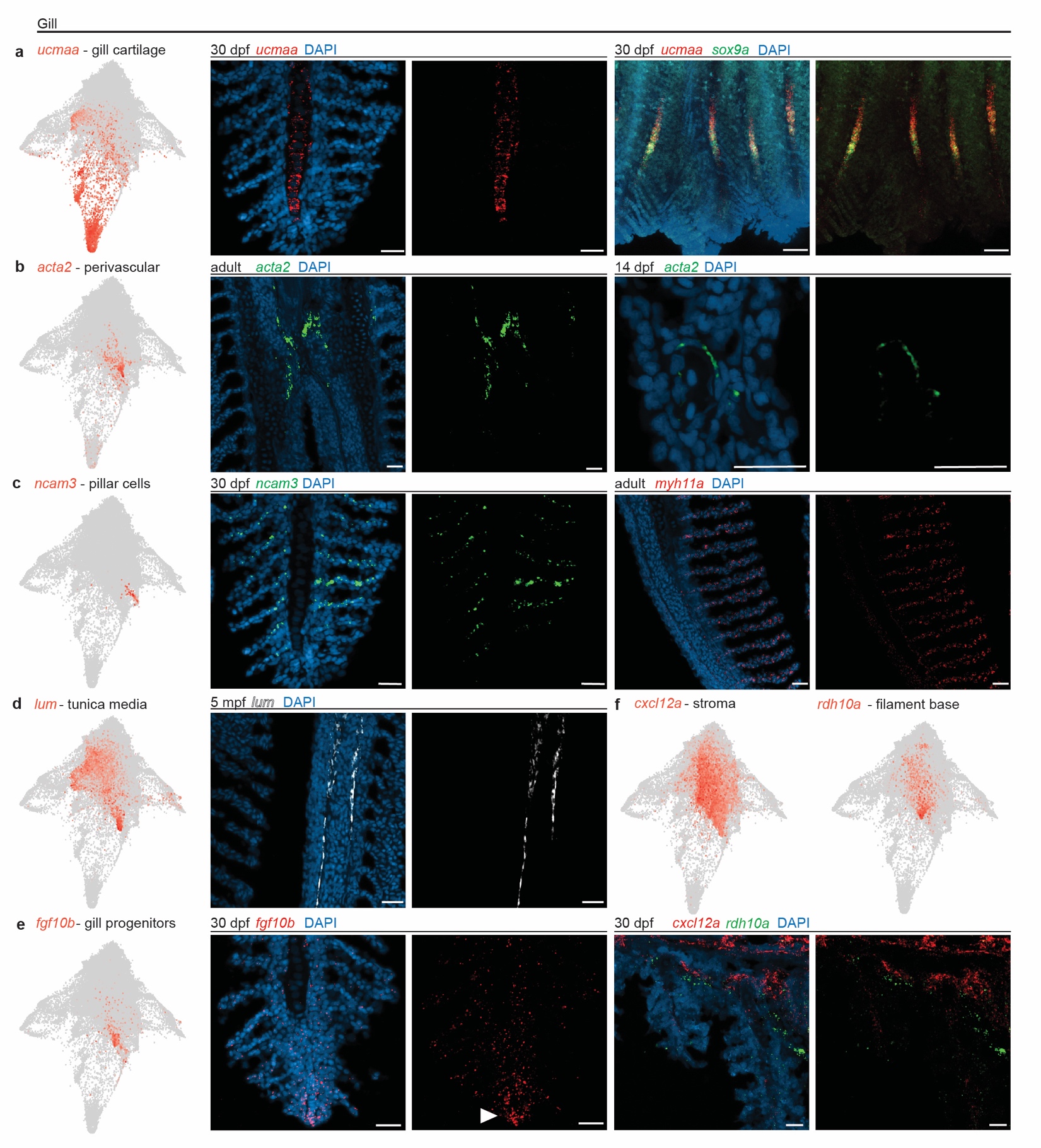
**Supplementary Figure 10. In vivo expression of markers for gill tissues. a-f**, STITCH plots and RNAscope in situ hybridization on sections of the gill filaments. At 30 dpf, *ucmaa* is expressed in cartilage of the primary filament, overlapping with the known cartilage marker *sox9a*. Perivascular expression of *acta2* is seen between primary filaments. At 14 dpf, *acta2* is seen around a large diameter blood vessel. Expression of *ncam3* is highly specific for pillar cells in the secondary filaments. Pillar cells are also labeled by *myh11a*. The tunica media surrounds the venous sinus in the primary filament and is labeled by *lum* expression. Expression of *fgf10b* marks progenitors at the tip (arrowhead) of the growing primary filament, in a reciprocal pattern to *rdh10a* at the base of the primary filament. DAPI labels nuclei in blue. Scale bars = 20 um.

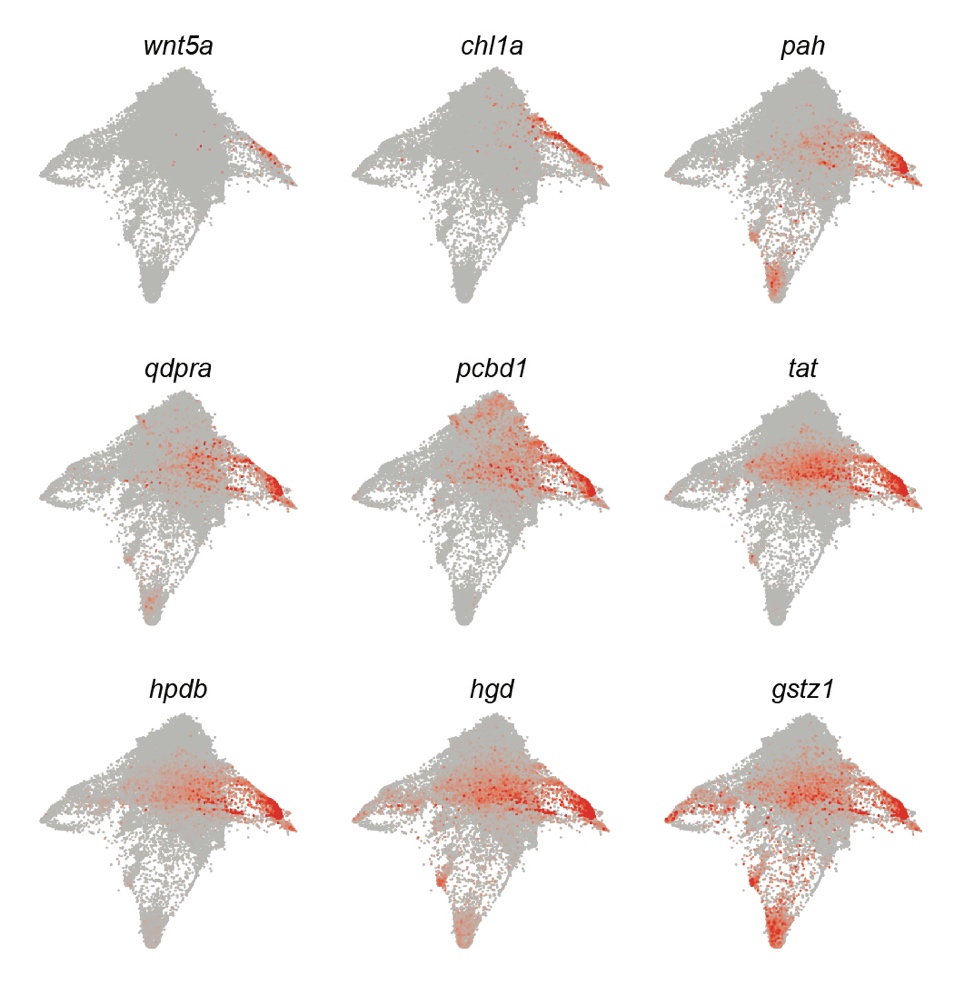

**Supplementary Figure 11. Dermal fibroblast markers.** STITCH plots expression of the cell adhesion molecule *chl1a* as early as 3 dpf in the branch leading to dermal fibroblasts. These specialized fibroblasts are also enriched for *wnt5a* and genes encoding all major enzymes of Phe/Tyr breakdown (*pah*, *qdpra*, *pcdb1*, *tat*, *hpdb*, *hgd*, *gstz1*).

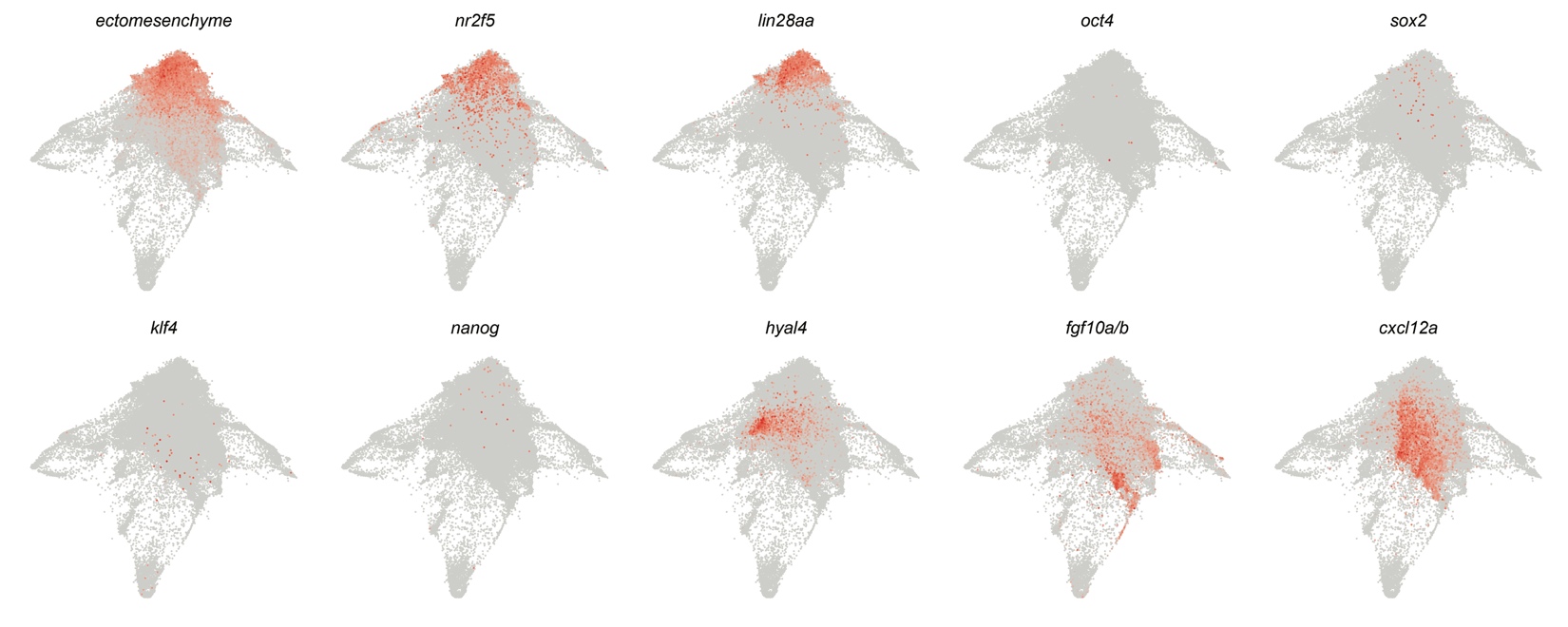
**Supplementary Figure 12. Transition from ectomesenchyme to region-specific progenitors.** STITCH plots show aggregated expression of 36 hpf ectomesenchyme genes, the ectomesenchyme-specific gene *nr2f5*, pluripotency-associated genes (*lin28aa*, *oct4/pou5f3*, *sox2*, *klf4*, *nanog*), perichondrium marker *hyal4*, gill progenitor markers *fgf10a* and *fgf10b* (summed), and stromal marker *cxcl12a*. The ectomesenchyme signature is rapidly extinguished and replaced by region-specific progenitor signatures (i.e. *hyal4*, *fgf10a/b*, *cxcl12a*), and we observe no evidence of a pluripotency signature.

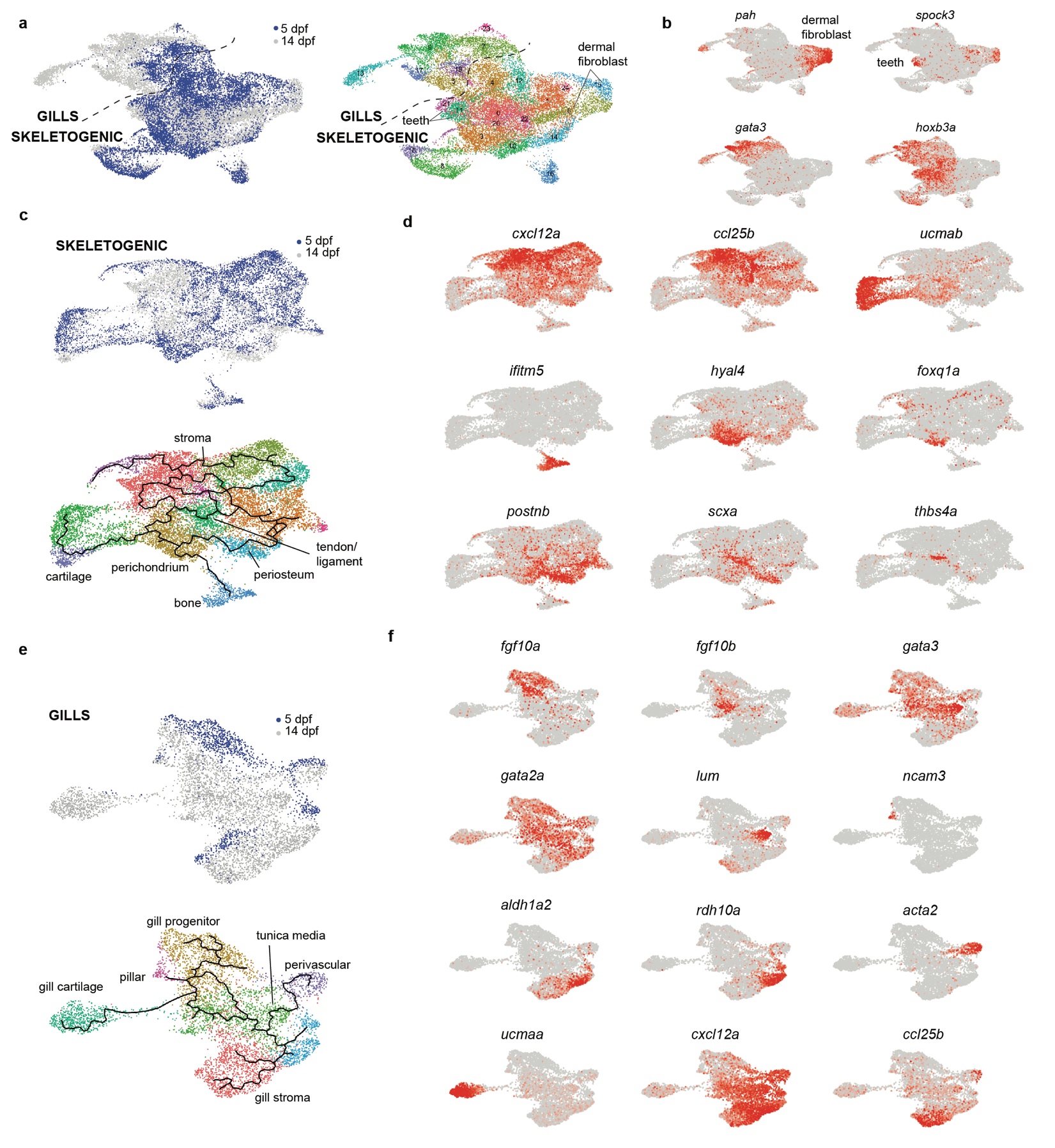

**Supplementary Figure 13. Pseudotime analysis of skeletogenic and gill subsets. a**, Combined 5 and 14 dpf scRNAseq datasets displaying cell types by stage and cluster (numbered), with dashed line demarcating skeletogenic versus gill subsets. **b**, Feature plots show genes used to remove dermal fibroblast (*pah*) and teeth (*spock3*) clusters. The gill subset was defined as positive for both *gata3* and *hoxb3a*. **c**, Skeletogenic sub-cluster displaying cell types by stage and cluster, with the lines showing pseudotime trajectories calculated by Monocle3. **d**, Feature plots for genes defining the labeled skeletogenic and stromal clusters. **e**, Gill sub-cluster displaying cell types by stage and cluster, with the lines showing pseudotime trajectories calculated by Monocle3. **f**, Feature plots for genes defining the labeled gill clusters.

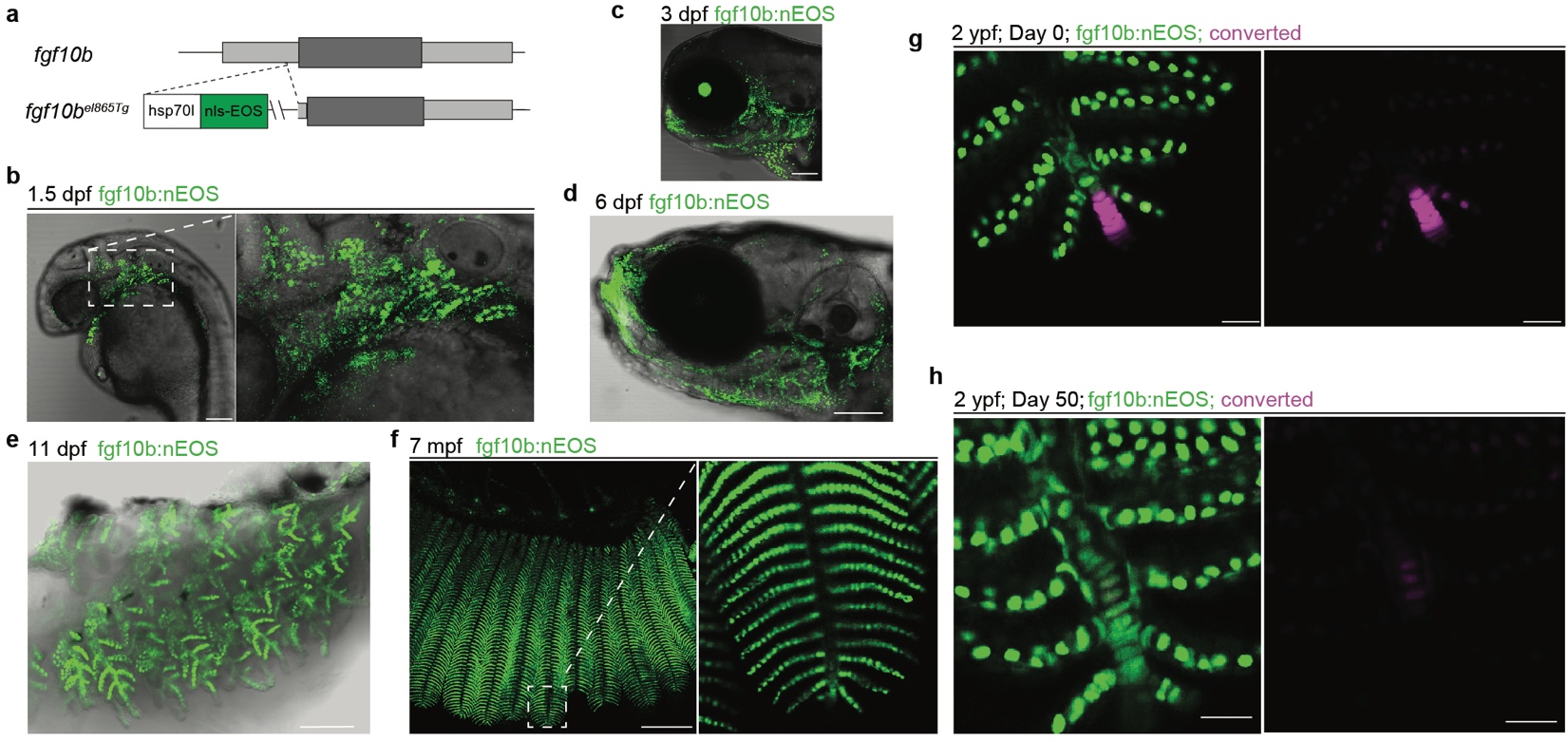

**Supplementary Figure 14. Characterization of *fgf10b:nls-EOS* fish. a**, Schematic showing insertion of the *hsp70l* minimal promoter followed by a fusion of the nuclear localization sequence and the EOS photoconvertible protein (nls-EOS) into the 5’ UTR region of the *fgf10b* gene locus. **b-d**. Confocal imaging of nEOS fluorescence and DIC light for context show expression in the gill region at 1.5 dpf and continuing through 3 and 6 dpf. Starting at 3 dpf, additional expression is seen in the frontonasal region. **e,f**, Confocal imaging of dissected gills show continued expression of *fgf10b:nls-EOS* in the filament system at 11 dpf and 7 months post-fertilization (mpf). **g,h**, In 2-year-old adults (*n*  = 2), UV-mediated photoconversion of *fgf10b:nls-EOS* from green to magenta at the tip of the primary filament resulted in labeling of new gill chondrocytes 50 days later. Scale bars = 100 um (a-e), 50 um (f), 20 um (g,h).

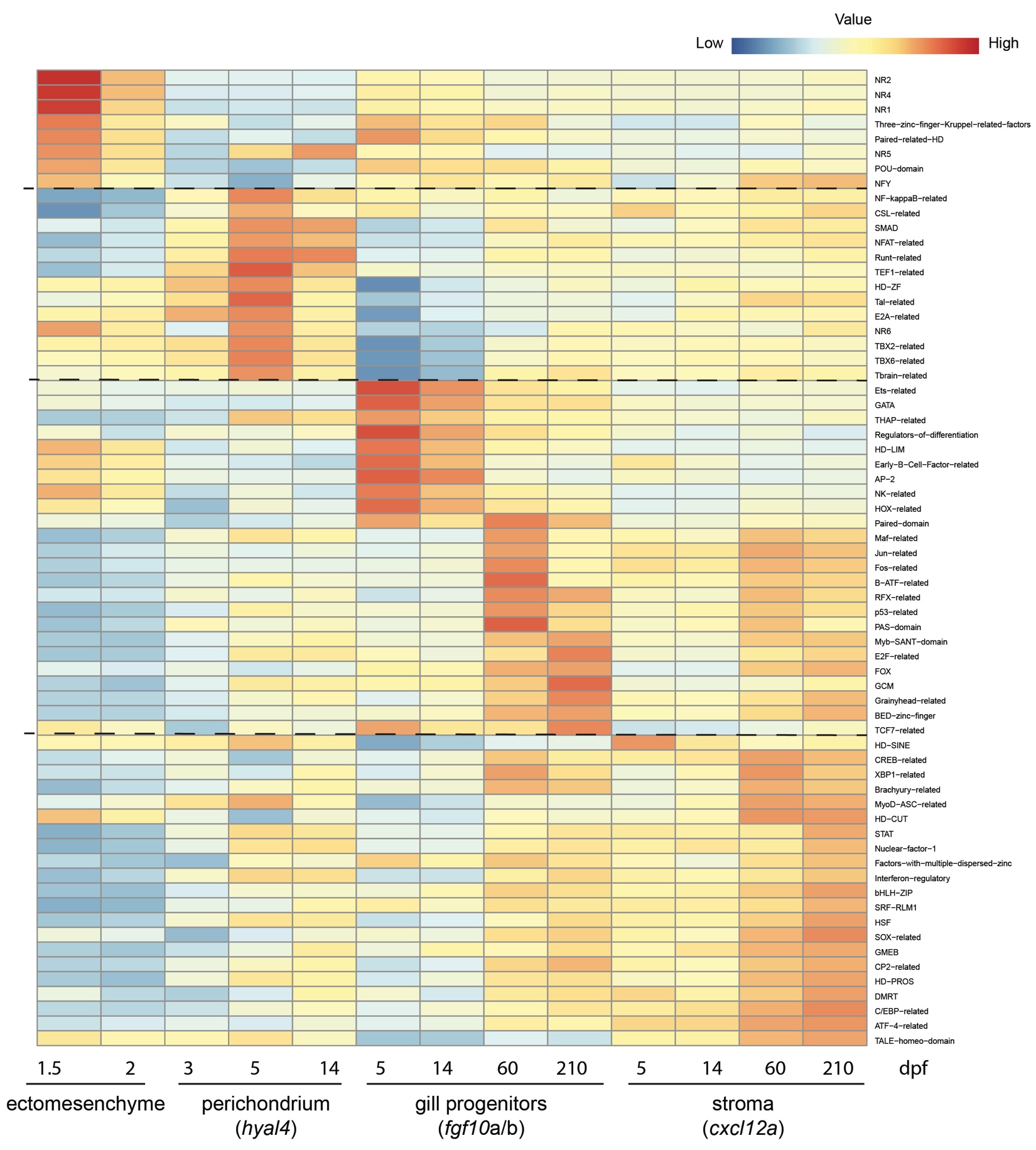

**Supplementary Figure 15. Motif enrichment in differentially accessible regions of mesenchyme cell types.** Shown are the relative enrichment of transcription factor binding motifs for the mesenchymal cell types at the indicated stages. Ectomesenchyme represents the aggregate of mesenchyme subsets at 1.5 and 2 dpf. The perichondrium clusters are defined by expression of *hyal4*, gill progenitor clusters by combined *fgf10a* and *fgf10b* expression*,* and stromal cell clusters by expression of *cxcl12a.*

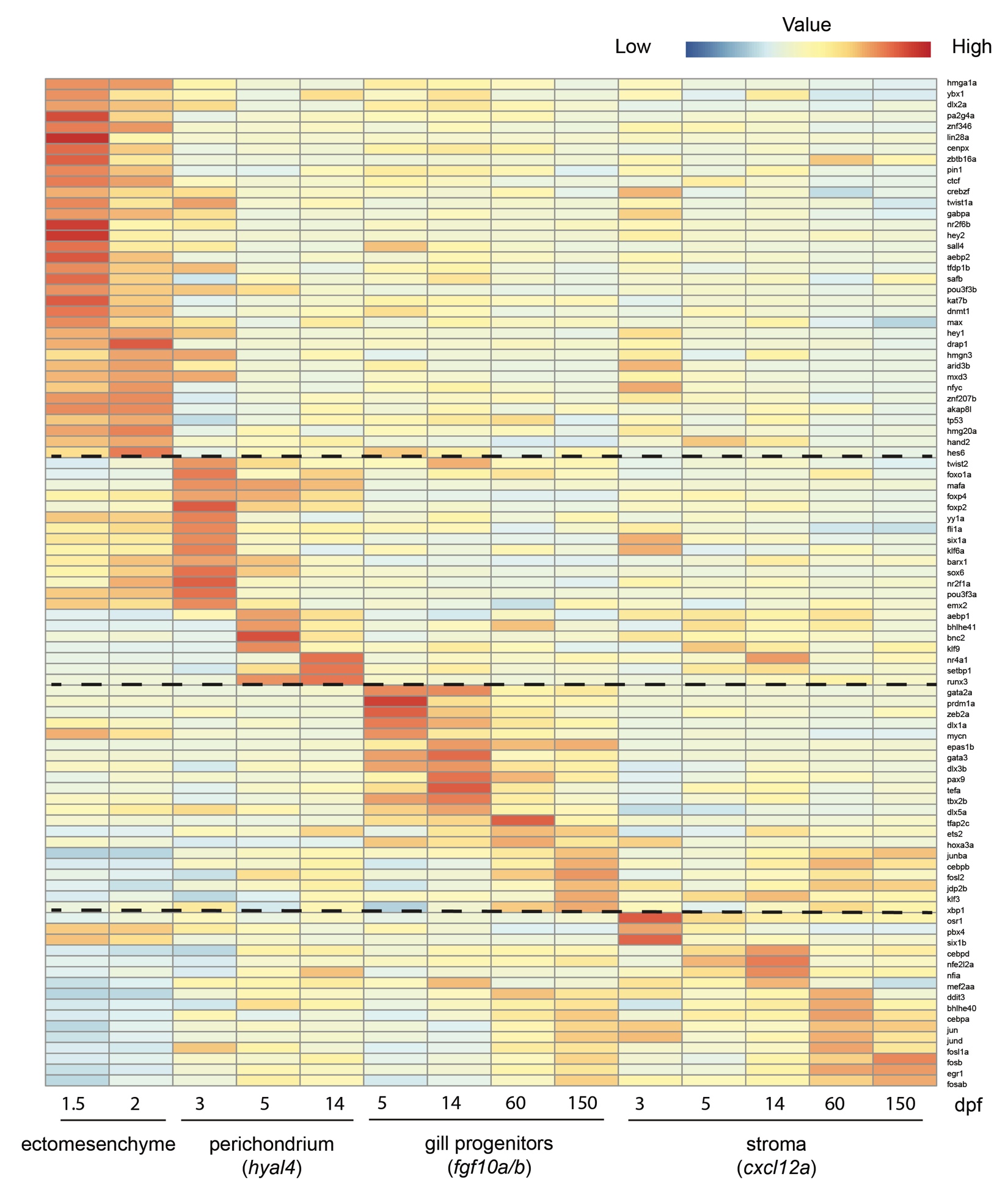
**Supplementary Figure 16. Transcription factor gene body activity enrichment in mesenchyme cell types.** Shown are the relative enrichment of transcription factor gene body activities (proxy for gene expression in snATACseq datasets) for the mesenchymal cell types at the indicated stages. Ectomesenchyme represents the aggregate of mesenchyme subsets at 1.5 and 2 dpf. The perichondrium clusters are defined by expression of *hyal4*, gill progenitor clusters by combined *fgf10a* and *fgf10b* expression*,* and stromal cell clusters by expression of *cxcl12a.*

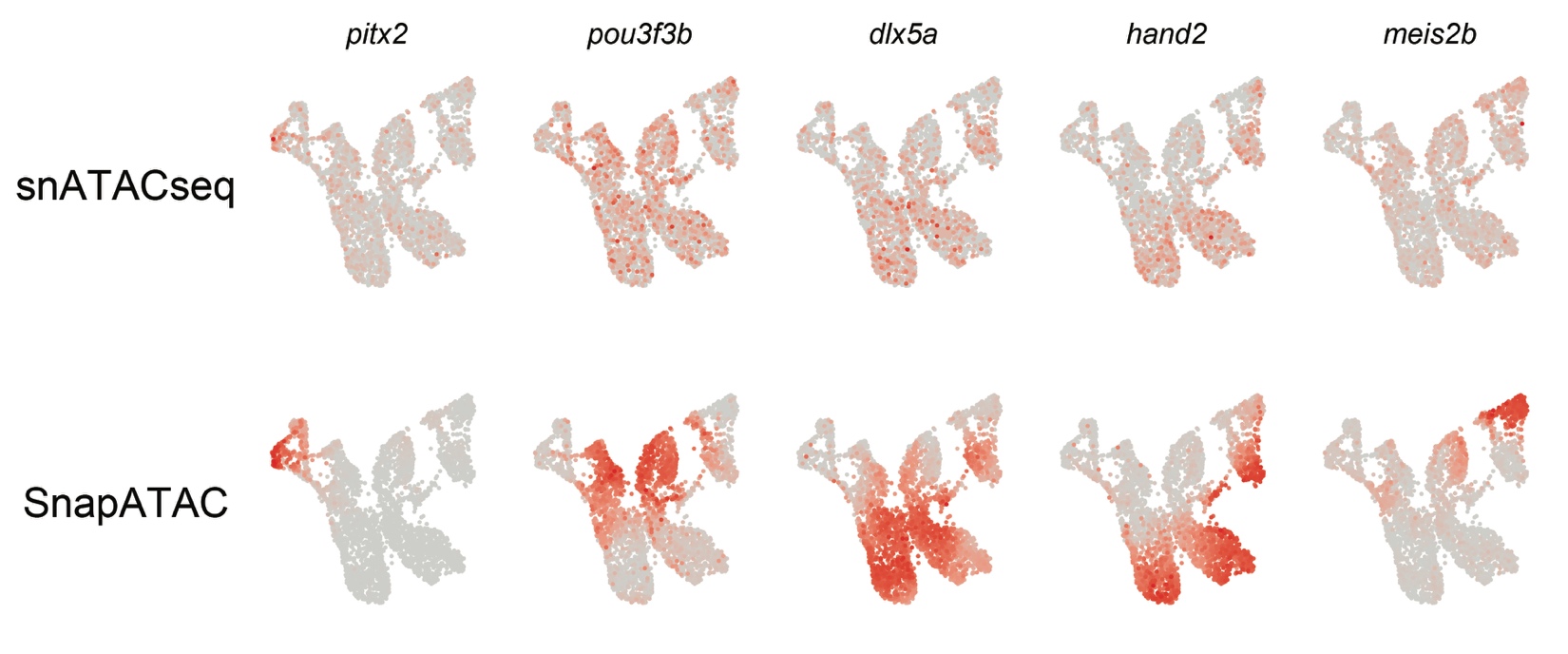
**Supplementary Figure 17. Comparison of SnapATAC to snATACseq for resolution of spatial expression at 1.5 dpf.** UMAPs for the indicated region-specific gene body activities as calculated by snATACseq data alone, versus integrated snATACseq and scRNAseq data (pseudo-multiome) using SnapATAC.

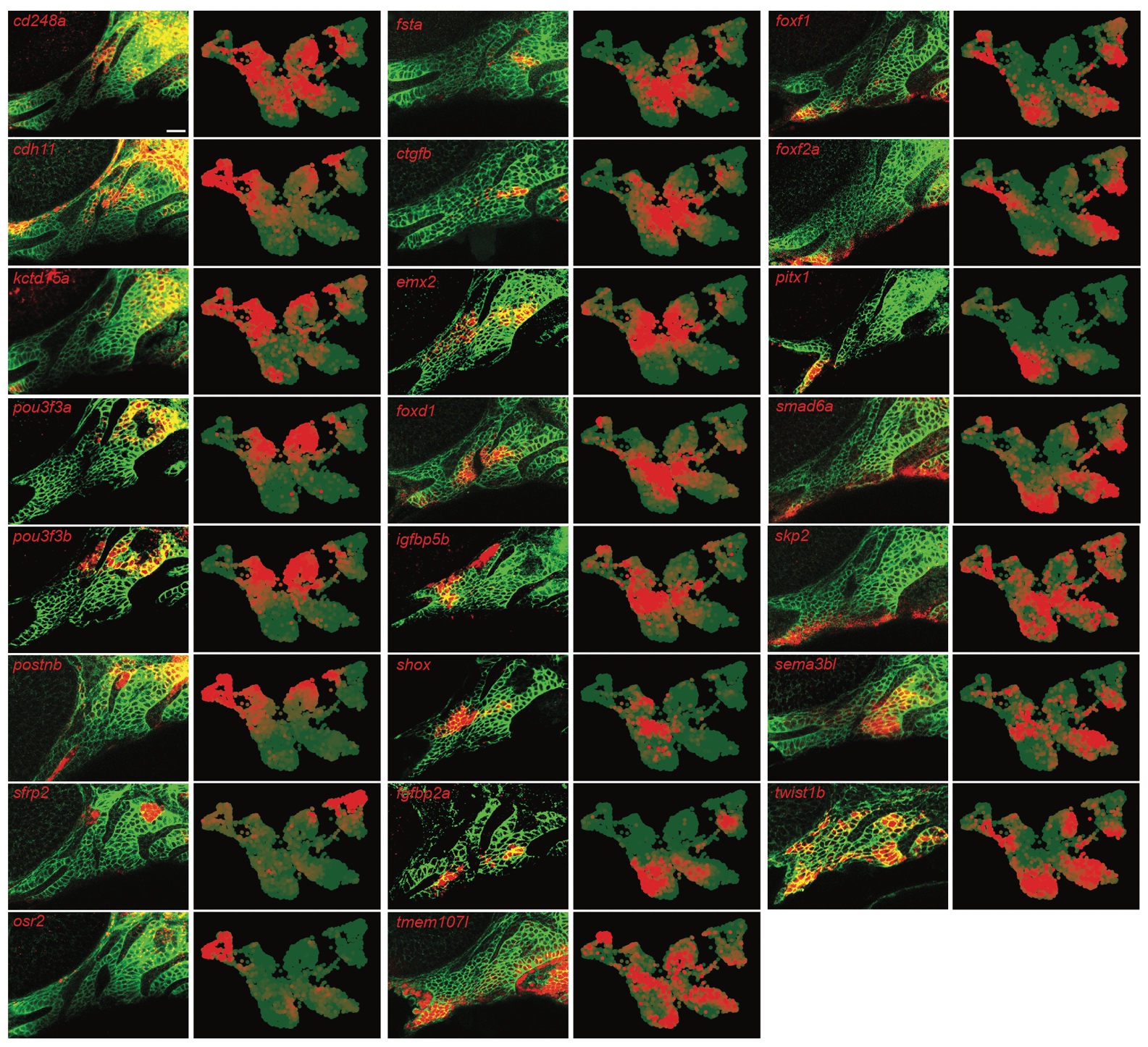

**Supplementary Figure 18. Comparison of SnapATAC spatial expression predictions to previously validated in vivo expression patterns.** For each gene, the left panel shows RNA in situ hybridization at 1.5 dpf (red) relative to *sox10:membrane-GFP*+ CNCCs of the mandibular and hyoid arches (green) as published in (Askary et al., 2017). Right panels show expression at 1.5 dpf as predicted by SnapATAC. Note that SnapATAC predicts expression for *sfrp2* in the posterior-most arches not imaged in (Askary et al., 2017), and for *osr2* expression in the frontonasal domain not imaged in (Askary et al., 2017).

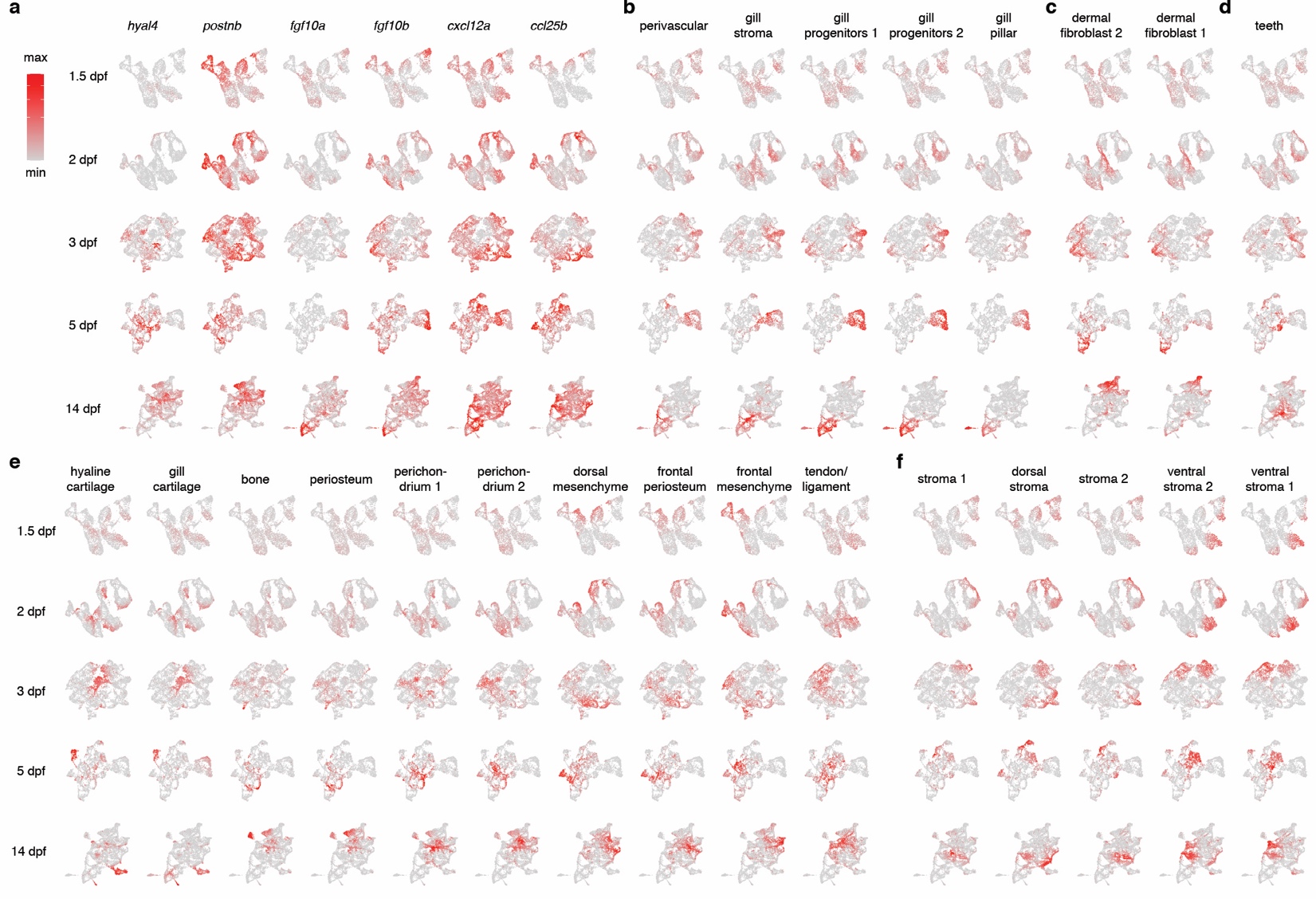

**Supplementary Figure 19. Retrograde mapping of peak module scores for each cluster at 14 dpf onto earlier stages. a**, UMAP feature plots show genes marking perichondrium (*hyal4*), periosteum (*postnb*), gill progenitors (*fgf10a* and *fgf10b*), and stromal cells (*cxcl12a* and *ccl25b*).**b-f**, For each of the 23 clusters at 14 dpf, peak module scores were calculated and projected onto UMAP plots across the indicated stages. The minimum cut-off for each peak module score is set to 0, and the heatmap was square root transformed to increase contrast. We grouped clusters for gill cell types (b), dermal fibroblasts (c), teeth (d), skeletogenic cells (e), and stromal cells (f).

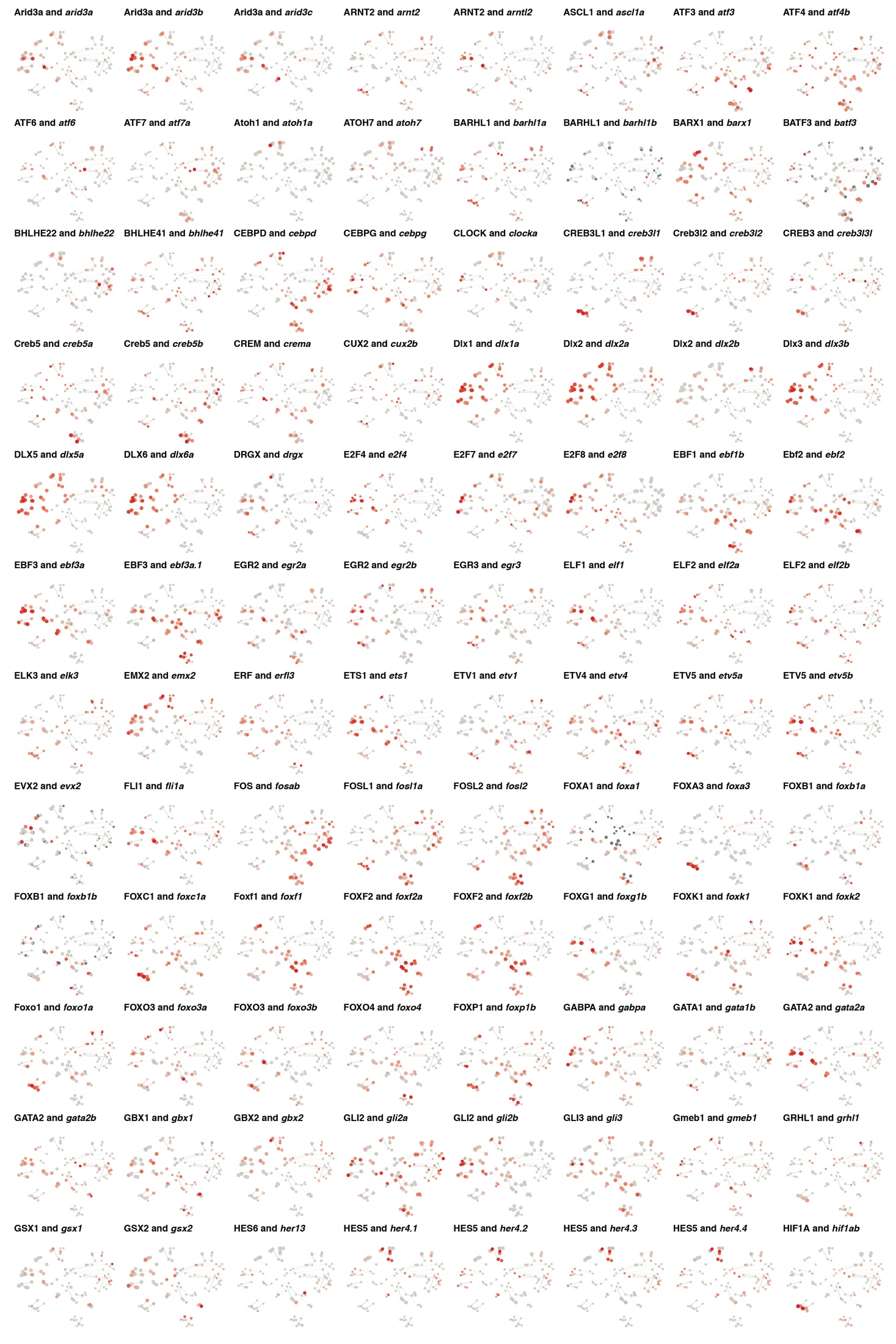

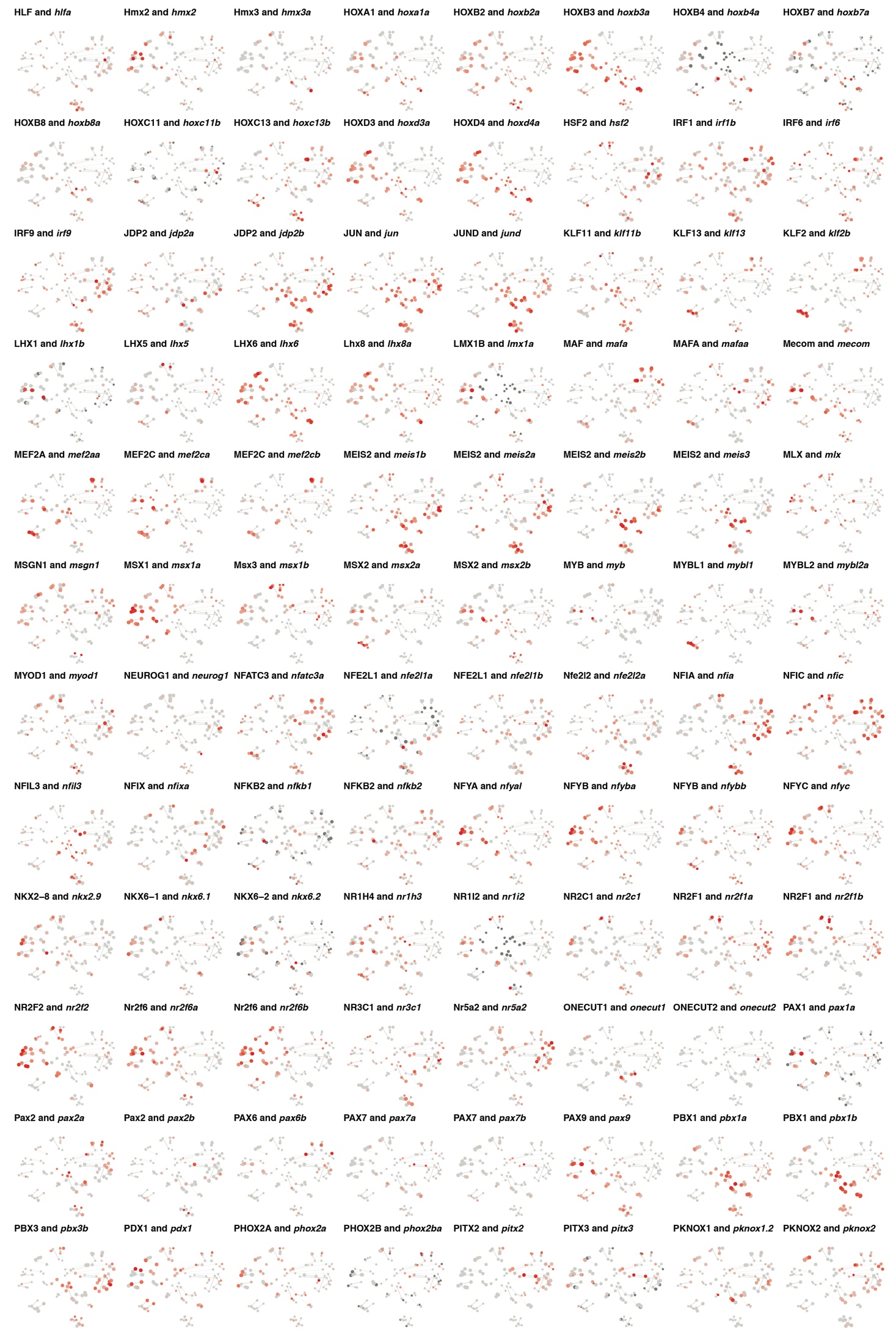

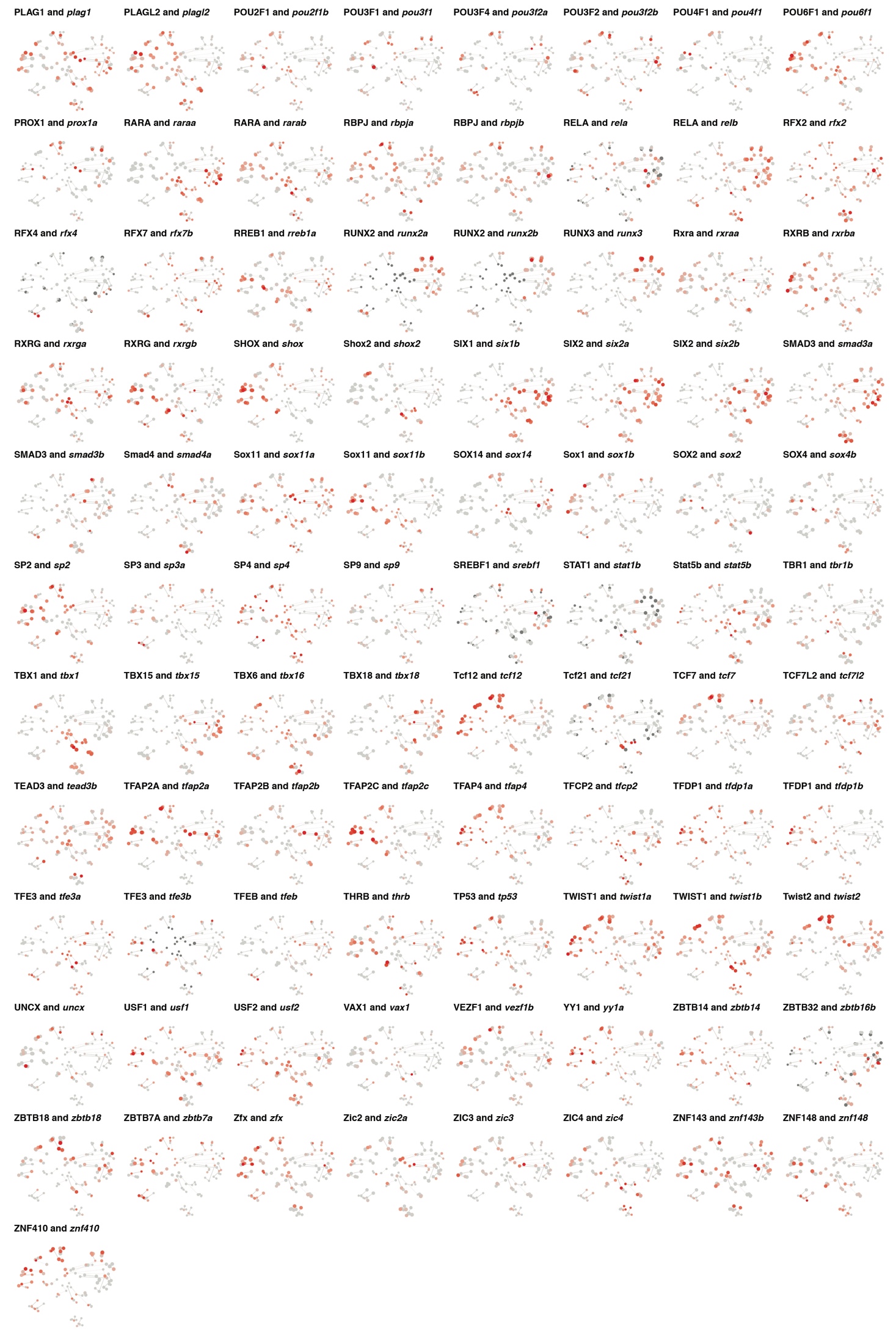
**Supplementary Figure 20 (previous 3 pages). Constellation plots of every motif and transcription factor pair.** Plotted is each transcription factor showing correlated gene body activity and binding motif enrichment in specific clusters of the Constellations map (see Figure 5 for more details). Size of circle shows correlation of motif enrichment with the cell cluster, and shade of red color indicates correlation of gene body activity (proxy of expression) with the cell cluster.

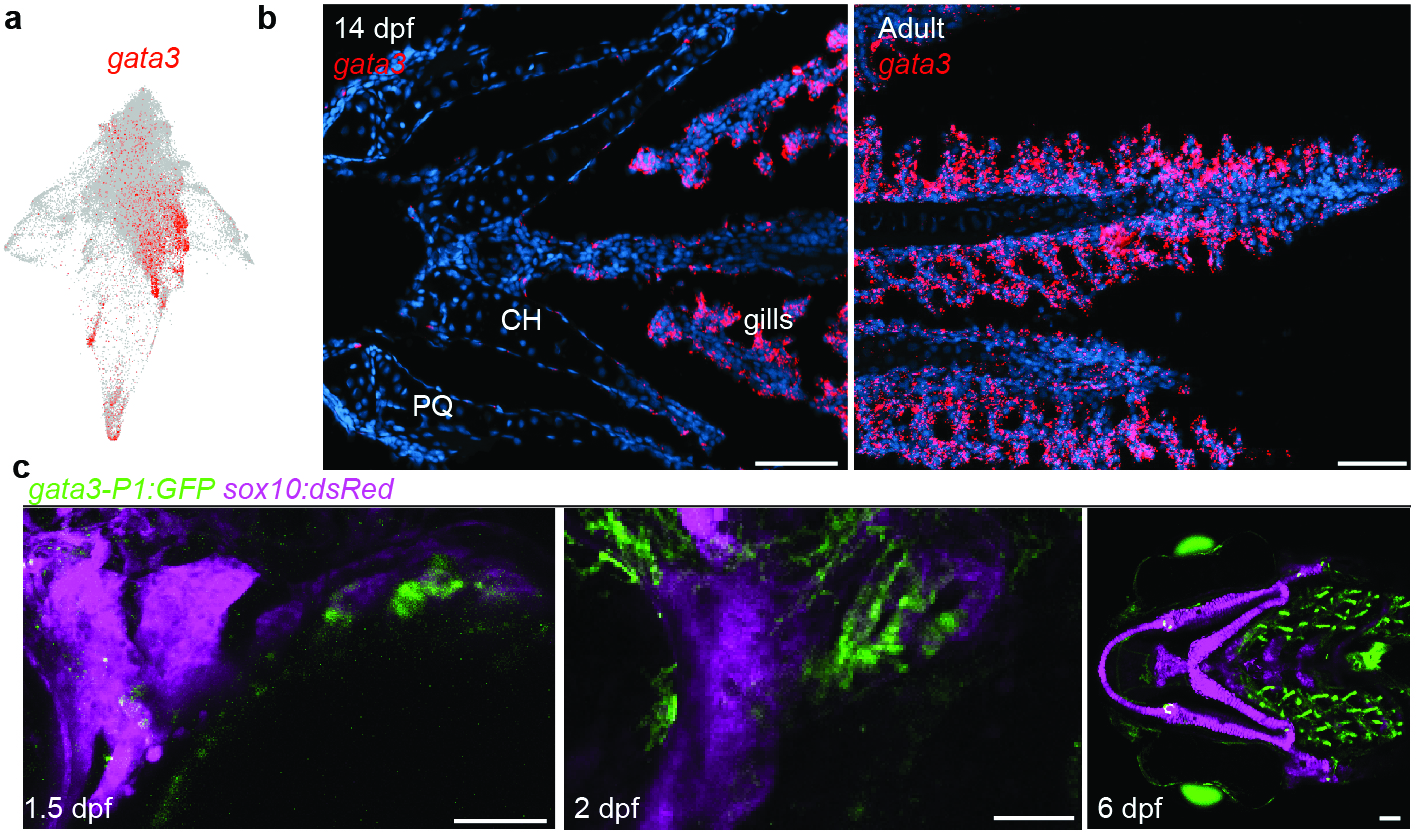

**Supplementary Figure 21. Gill-specific expression of *gata3* and the *gata3-P1:GFP* transgenic line. a**, STITCH feature plot shows enrichment of *gata3* expression in gill-associated populations. **b**. RNAscope in situ hybridizations for *gata3* show selective expression in the forming gills at 14 dpf and continued gill filament expression in 2-year-old adult fish. Note absence of *gata3* expression in the more anterior ceratohyal (CH) and palatoquadrate (PQ) cartilages. DAPI labels nuclei in blue. **c**, *gata3-P1:GFP* drives expression in the gill-forming posterior arches at 1.5 and 2 dpf and the developing gill filaments at 6 dpf. For reference, *sox10:dsRed* labels arch CNCCs at 1.5 and 2 dpf and cartilage at 6 dpf. Scale bars = 50 um.

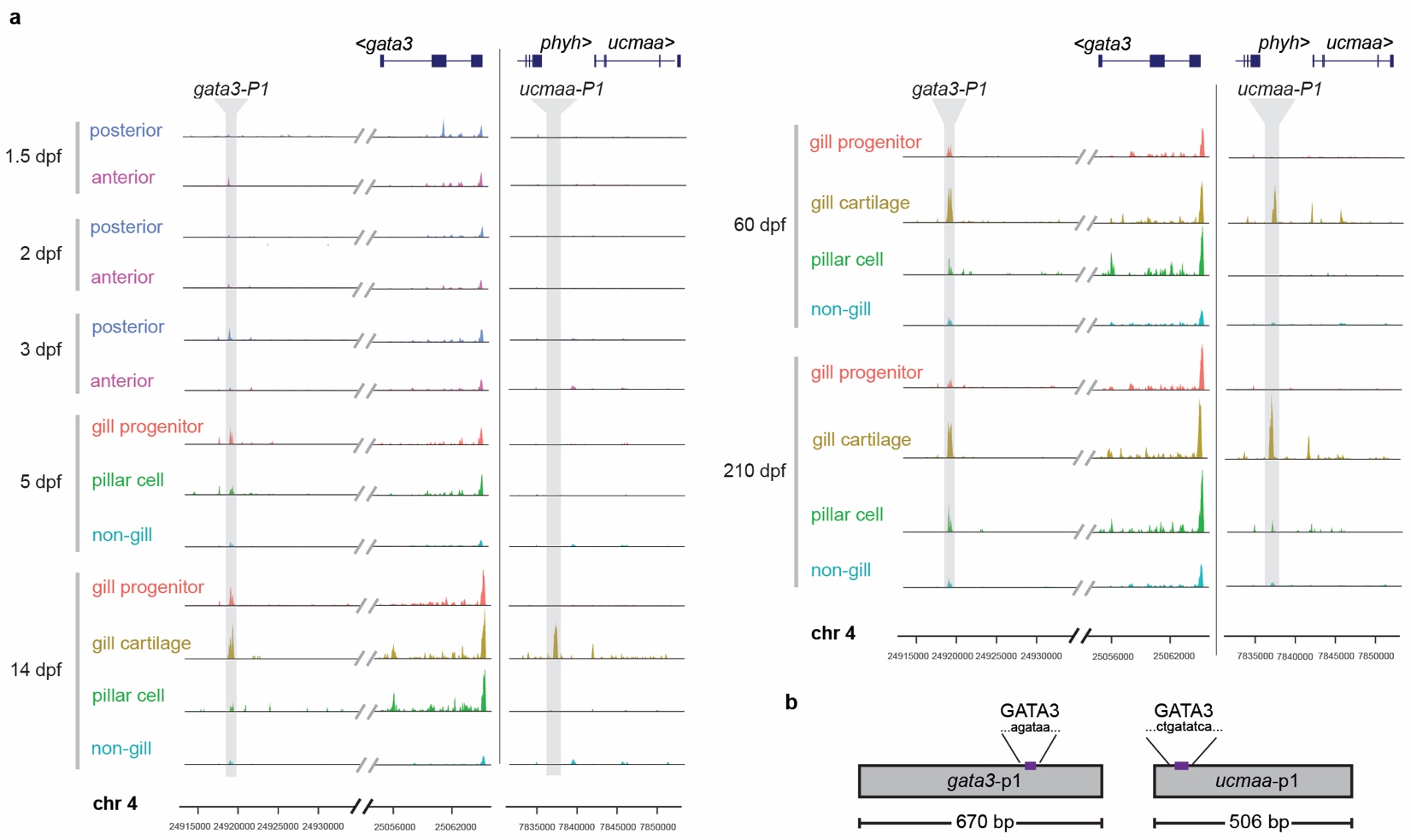

**Supplementary Figure 22. Gill-specific *gata3* and *ucmaa* enhancers. a**, Genomic tracks for aggregated posterior arch and anterior arch clusters at 1.5, 2, and 3 dpf, and gill progenitor, gill pillar cell, gill cartilage, and aggregated non-gill clusters at later stages. The y-axis displays normalized chromatin accessibility (range 0-240) in snATACseq data across the *gata3* and *ucmaa* loci on chromosome 4 (genomic coordinates at bottom correspond to GRCz11 zebrafish genome assembly). The validated gata3-P1 and ucmaa-P1 gill-specific enhancers are shown in grey. **b**, Predicted GATA3 binding sites are shown for each enhancer.

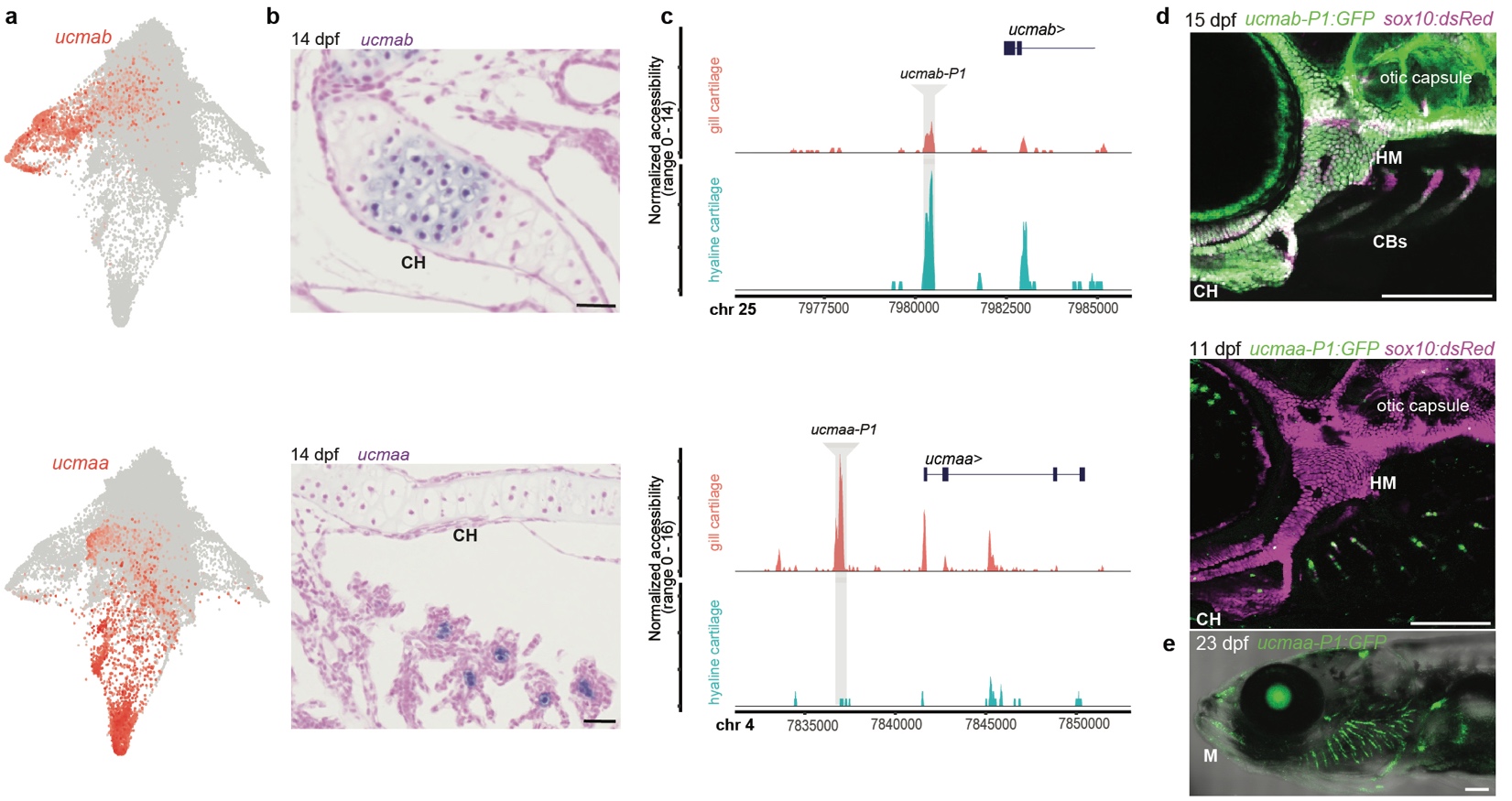

**Supplementary Figure 23. Comparison of hyaline and gill cartilage.** **a**. STITCH feature plots show selective expression of *ucmab* in the hyaline cartilage branch and *ucmaa* in the gill cartilage branch. **b**, Colorimetric RNA in situ hybridization on facial sections show *ucmab* expression in the ceratohyal growth plate and *ucmaa* expression in gill cartilage (blue is expression and nuclear fast red counterstain shows tissue context). **c**, Genome tracks of 60 dpf snATACseq data show selective accessibility of a ucmab-P1 peak in hyaline cartilage and a ucmaa-P1 peak in gill cartilage. **d**, Confocal imaging shows expression of *ucmab-P1:GFP* in hyaline cartilage of the hyomandibula (HM), ceratohyal (CH), ceratobranchials (CBs), and otic capsule, and *ucmaa-P1:GFP* in gill filament cartilage. *sox10:dsRed* labels all cartilage. **e**, Confocal imaging at 23 dpf shows expression of *ucmaa-P1:GFP* in gill filament cartilages, as well as some expression in the permanent Meckel’s (M) cartilage. DIC channel shows tissue context in white. Scale bars = 20 um (b), 200 um (d,e).

**Supplementary Table 1. Cluster marker genes, gene body activities, and motifs for each single-cell experiment.** Shown are the top marker genes for each scRNAseq dataset, and top gene body activities and motifs for each snATACseq dataset. Significant cluster markers for zebrafish craniofacial cell types derived from scRNAseq and snATACseq (*P* value less than 0.001). Sheets with datasets are organized chronologically (1.5, 2, 3, 5, 14, 60, 150, 210 dpf). Every sheet has UMAP visualization with the unsupervised clustering.

**Supplementary Table 2. Gene ontology, motif family, and TF analysis of mesenchyme population in scRNAseq data. Sheet 1,** Gene ontology analysis of biological process (BP) for each mesenchymal cluster. **Sheet 2,** Scaled means of every motif family of each mesenchymal cluster. **Sheet 3,** Scaled means of every TF of each mesenchymal cluster.

**Supplementary Table 3. Peaks used for tissue module score calculations in the Constellations analysis.** Genomic coordinates (GRCz11 build) are shown for the top peaks used to calculate module scores for each of the 23 clusters identified at 14 dpf.

**Supplementary Table 4. Top correlated motifs and TFs to tissue module scores at each skewed stage.** List of top 20 TFs with highest coefficients for each tissue module score at each skewed time point. The list of these top TFs is used to search for their corresponding motifs which exist in the top 100 highest coefficients for each tissue module score at each skewed time point. Lists of motifs and TFs are in descending order by their degree of correlation.
